# NSUN5 is essential for proper Hippo signaling in mouse preimplantation development

**DOI:** 10.1101/2023.09.08.556795

**Authors:** Dan Liu, Takuto Yamamoto, Naojiro Minami, Shinnosuke Honda, Shuntaro Ikeda

## Abstract

**Abstract:** NOL1/NOP2/Sun domain family, member 5 (NSUN5) is an enzyme belonging to the 5-methylcytosine (m5C) writer family that modifies rRNA and mRNA. By re-analyzing published RNA-sequencing data, we found that *Nsun5* expression was increased at the 2-cell stage during mouse preimplantation development. It has been reported that m5C regulates the stability of rRNA and mRNA. Therefore, NSUN5 may affect the expression of genes required for the development and differentiation of early mouse embryos via its role in modifying RNA. The Hippo signaling pathway has been identified as a critical regulator of the segregation of inner cell mass and trophectoderm lineages during mouse embryogenesis. According to these findings, we hypothesized that NSUN5 controls cell differentiation by regulating the expression of components of the Hippo signaling pathway in mouse early embryos. Using *Nsun5*-specific small interfering RNAs, we efficiently knocked down *Nsun5* expression in mouse preimplantation embryos, resulting in impairments in early development, including reduced blastocyst formation, smaller size, and impaired hatching from the zona pellucida. *Nsun5* depletion also led to decreased cell numbers, suggesting impaired cell proliferation. Furthermore, *Nsun5* knockdown embryos exhibited reduced yes-associated protein 1 (YAP1) nuclear translocation during the morula stage, potentially affecting cell differentiation. The imbalance in pluripotent and trophectoderm lineages was evident with a significant reduction in the ratio of CDX2-positive cells to OCT4-positive cells. *Nsun5* depletion impacted the Hippo signaling pathway, increasing the expression of the key genes *Lats1* and *Lats2* during the morula stage. Our findings underscore the essential role of *Nsun5* in early embryonic development by affecting cell proliferation, YAP1 nuclear translocation, and the Hippo pathway. Understanding the roles of *Nsun5* in mouse early embryos will provide insights into complex regulatory networks in development, with implications for reproductive medicine and fertility research.

## Introduction

The dynamic modification of RNA molecules, such as rRNA and mRNA, plays a critical role in gene expression regulation (Roundtree, et al. 2017). Specifically, 5-methylcytosine (m5C) modifications are associated with the stability of rRNA and mRNA (Gao and Fang 2021). NOL1/NOP2/Sun domain family, member 5 (NSUN5), a member of the m5C writer family, is known for its role in modifying rRNA and mRNA. Previous studies have demonstrated that *Nsun5* knockout mice exhibit impaired follicular development, ovarian function, and embryonic development (Yang, et al. 2019).

The Hippo signaling pathway is a crucial regulator of the segregation of inner cell mass and trophectoderm lineages during mouse embryogenesis (Karasek, et al. 2020, Mihajlović and Bruce 2016). Activation of the Hippo pathway is tightly controlled by molecular mechanisms, including apical cell polarity complexes (PAR3/PAR6/aPKC) and large tumor suppressor 1/2 (LATS1/2) (Hirate, et al. 2015). The downstream effector protein yes-associated protein 1 (YAP1) plays a redundant role in cellular responses (Nishioka, et al. 2009b, Rayon, et al. 2014). The activated LATS1/2 phosphorylate YAP1 and inhibit its nuclear translocation and binding to TEA domain family member 4 (TEAD4), which is required for the specific activation of trophectoderm-associated genes, such as caudal type homeobox 2 (*Cdx2*) (Hirate, et al. 2015, Rayon, et al. 2014). The activation of LATS1/2 relies on the presence of MOB kinase activator 1A and 1B (MOB1A/B) (Bothos, et al. 2005). The pivotal role of the Hippo signaling pathway in embryonic development is evident from studies demonstrating that deletion of *Mob1a*/*b* in mice results in early embryonic lethality during the preimplantation stage (Nishio, et al. 2012).

Furthermore, the Notch signaling pathway has been identified as a key regulator of embryonic development, with its blockade leading to the downregulation of *Sox2* transcription and delayed embryo hatching (Batista, et al. 2020). Recent findings have highlighted an interplay between the Notch and Hippo signaling pathways (Liu, et al. 2019). YAP1/TEAD transcription factors directly regulate genes in both pathways, including trophoblast genes (*Cdx2*, *Eomes*, and *Gata3*) (Nishioka, et al. 2009a, Ralston, et al. 2010, Yimlamai, et al. 2014) and *Ccnd1* (Mizuno, et al. 2012).

Previous studies have shown m5C modifications in the 3′ untranslated region of *Yap1*, which increase the stability of *Yap1* mRNA (Yu, et al. 2023). Given these intriguing findings, we hypothesized that *Nsun5* may play a regulatory role in cell differentiation in mouse early embryos by modulating the expression of key components of the Hippo signaling pathway. To test our hypothesis, we employed an experimental approach involving microinjection of small interfering RNA (siRNA) targeting *Nsun5* into fertilized zygotes. By analyzing the resulting embryos at the morula and blastocyst stages and using specific antibodies for immunostaining (YAP1 for morulae; CDX2 and OCT4 for blastocysts), we aimed to determine the impact of *Nsun5* knockdown on the cell differentiation of mouse preimplantation embryos.

## Materials and Methods

### Re-analysis of published RNA-sequencing (RNA-seq) data

To investigate *Nsun5* expression levels during mouse preimplantation development, we analyzed the publicly available RNA-seq data from a recent study (GSE71434) (Zhang, et al. 2016). The fastq files (2 replicates) for MII oocytes; 1-cell, early 2-cell, late 2-cell, 4-cell, and 8-cell stage embryos; and inner cell mass of blastocysts were quality-controlled and mapped to the mouse genome (mm10) using STAR (v. 2.7.10a) (Dobin, et al. 2013). The mapped reads were fed to RSEM (v. 1.3.3) (Li and Dewey 2011) to calculate the transcripts per million values for *Nsun5*.

### Collection of oocytes, *in vitro* fertilization, and embryo culture

ICR female mice (8–12 weeks old; Japan SLC, Shizuoka, Japan) were superovulated by intraperitoneal injection of 7.5 IU equine chorionic gonadotropin (ASUKA Animal Health, Tokyo, Japan) followed 46–48 h later by 7.5 IU human chorionic gonadotropin (ASUKA Animal Health). Cumulus-oocyte complexes were collected from the ampullae of the excised oviducts at 14 h after human chorionic gonadotropin injection. Cumulus-oocyte complexes were placed in a 100-μL droplet of human tubal fluid medium supplemented with 4 mg/mL bovine serum albumin (BSA; A3311; Sigma Aldrich, St. Louis, MO) (Minami, et al. 2001). Spermatozoa were collected from the cauda epididymis of 12–18-week-old ICR male mice (Japan SLC) and cultured for at least 1 h in a 100-μL droplet of human tubal fluid medium. After preincubation, the sperm suspension was added to fertilization droplets at a final concentration of 1.0 × 10^6^ cells/mL. At 2 h post-insemination (hpi), morphologically normal fertilized embryos were collected and cultured in potassium simplex optimized medium supplemented with amino acids (Ho, et al. 1995) and 1 mg/mL BSA under paraffin oil (Nacalai Tesque, Kyoto, Japan). All incubations were performed at 37°C under 5% CO2 in air.

### Microinjection of siRNA

siRNA molecules were microinjected into the cytoplasm of zygotes (1-cell embryos) under an inverted microscope (IX73; Olympus, Tokyo, Japan) equipped with a piezo injector (PMAS-CT150; PRIME TECH, Tokyo, Japan) and a micromanipulator (IM-11-2; Narishige, Tokyo, Japan). All fertilized zygotes were microinjected with either non-targeting (control) or *Nsun5*-targeting siRNA (knockdown) within 3–6 hpi. Approximately, 10 pL siRNA (20 µM) solution was injected into the cytoplasm of each zygote. The siRNA sequences were as follows: si*Nsun5*-1s, CCUUUCCAGGGUCUGAACAtt; si*Nsun5*-1as, UGUUCAGACCCUGGAAAGGtt; si*Nsun5*-2s, GCAAUGGAUCCAGAACCUCtt; si*Nsun5-*2as, GAGGUUCUGGAUCCAUUGCtt.

### RNA extraction and reverse transcription-quantitative PCR (RT-qPCR)

Total RNA extraction and cDNA synthesis from embryos were performed using a SuperPrep^TM^ II Cell Lysis & RT Kit for qPCR (TOYOBO, Osaka, Japan). Synthesized cDNA was mixed with specific primers and KOD SYBR qPCR Mix (TOYOBO), followed by RT-qPCR amplification. The protocols for RT-qPCR and measurement of transcript levels were performed as described previously (Shikata, et al. 2020), and *H2afz* was used as an internal control. Relative gene expression was calculated using the 2−ΔΔCt method (Livak and Schmittgen 2001). The following primer sequences were used for RT-qPCR: *H2afz*, 5′-GGTAAAGCGTATCACCCCTCG-3′ (forward) and 5′-CTTCCCGATCAGCGATTTGTG-3′ (reverse); *Nsun5*, 5′-TGTATTCCAGCAACTTCCAGAACC-3′ (forward) and 5′-AAGGTGCGGTCTCAGCTTCTT-3′ (reverse); *Lats1*, 5′-AGGCGGATGTAGGAAGACCT-3′ (forward) and 5′-TCCATTGCTTGGGTGAGCTT-3′ (reverse); *Lats2*, 5′-AAATGAGAGCCACCCCGAAG-3′ (forward) and 5′-ACATCCCGCATTCACCAACT-3′ (reverse); *Sav1*, 5′-GGATGCTGTCCCGCAAGAAA-3′ (forward) and 5′-AAGGCATGAGATTCCGCAGC-3′ (reverse); *Mob1a*, 5′-CAGCAGCCGCTCTTCG-3′ (forward) and 5′-CCTCAGATTGCCACTTCCGA-3′ (reverse); *Mob1b*, 5′-CGACATGAGCTTCTTGTTTGGT-3′ (forward) and 5′-TGTGGCTTCCGCATGCTTTA-3′ (reverse); *Map4k4*, 5′-GTGACTTCCGTGGTGGGATT-3′ (forward) and 5′-TTGACCACTGAGCCTTTCCG-3′ (reverse); *Nf2*, 5′-TGCTGTCCAGGCCAAGTATG-3′ (forward) and 5′-CCTCCCACATTTCCGGAGTC-3′ (reverse); *Notch2*, 5′-CCGTGGGGCTGAAAAATCTC-3′ (forward) and 5′-GGGTCATCTTCCGACAGCAA-3′ (reverse); *Notch1*, 5′-CACCAGGGTGGTCAGGAAAA-3′ (forward) and 5′-GGGCAGCGACAGATGTATGA-3′ (reverse); *Hes1*, 5′-GAGGCTGCCAAGGTTTTTGG-3′ (forward) and 5′-ACTTTACGGGTAGCAGTGGC-3′ (reverse); Sox2, 5′-AAACCACCAATCCCATCCA-3′ (forward) and 5′-CCCCAAAAAGAAGTCCCAAG-3′ (reverse); Cdx2, 5′-AGCTGCTGTAGGCGGAATGTATG-3′ (forward) and 5′-TCAGTGACTCGAACAGCAGCAA-3′ (reverse); *Gata3*, 5′-CCAAGCGAAGGCTGTCGG-3′ (forward) and 5′-GTCCCCATTAGCGTTCCTCC-3′ (reverse).

### Immunofluorescence analysis

Embryos were fixed in 4% paraformaldehyde in phosphate-buffered saline (PBS) at room temperature for 20 min after removal of the zona pellucida with acid Tyrode’s solution (pH 2.5). After washing three times in PBS containing 0.3% polyvinylpyrrolidone (K-30; Nacalai Tesque), fixed embryos were treated with 0.5% Triton X-100 (Sigma-Aldrich) in PBS at room temperature for 40 min, and were blocked in PBST containing 1.5% BSA, 0.2% sodium azide, and 0.02% Tween 20 (antibody dilution buffer) at room temperature for 1 h. Embryos were incubated with one of the following primary antibodies in antibody dilution buffer at 4°C overnight: anti-YAP1 (1:2,000 dilution; DBH1X; Cell Signaling, Danvers, MA), anti-CDX2 (1:100 dilution; MU392A-UC; Biogenex, Fremont, CA), and anti-OCT4 (1:200 dilution; ab19857; Abcam, Cambridge, UK). Embryos were washed three times in antibody dilution buffer and then incubated with an appropriate secondary antibody in antibody dilution buffer (1:500 dilution; Alexa Fluor 488-conjugated goat anti-rabbit IgG or Alexa Fluor 594-conjugated goat anti-mouse IgG; Invitrogen) at room temperature for 1 h. After staining with Hoechst 33342 (Sigma-Aldrich), embryos were mounted on slides in Vectashield mounting medium (Vector Laboratories, Newark, CA) and signals were observed using a fluorescence microscope (BX53 or IX73; Olympus, Tokyo, Japan). The number of analyzed embryos is given in each figure legend.

### Statistical analysis

Developmental rates were analyzed using one-way analysis of variance (ANOVA) followed by the Tukey-Kramer test for multiple comparisons. For gene expression analysis, RT-qPCR data were analyzed using Student’s *t*-test for pairwise comparisons or one-way ANOVA followed by the Tukey-Kramer test for multiple comparisons. P-values < 0.05 were considered statistically significant; 0.05 < P < 0.1 were considered a tendency toward significance.

### Ethical approval for the use of animals

All experimental procedures were approved by the Animal Research Committee of Kyoto University (permit no. R3-17) and performed in accordance with the committee’s guidelines. Mice were euthanized by cervical dislocation without the usage of anesthetic agents by trained personnel following the ARRIVE guidelines.

## Results

### siRNA efficiently knocked down the *Nsun5* expression

Our re-analysis of published RNA-seq data revealed a significant increase in *Nsun5* expression at the 2-cell stage during mouse preimplantation development (Fig. 1a). To efficiently knock down *Nsun5* and minimize off-target effects, we employed two specific siRNAs (si*Nsun5*-1 and si*Nsun5*-2) to target its mRNA. Embryos were collected at 24 h after microinjection for RT-qPCR analysis to assess knockdown efficiency. The qPCR results demonstrated that both siRNAs exhibited good efficiency in knocking down *Nsun5* levels (Fig. 1b).

**Fig. 1.**
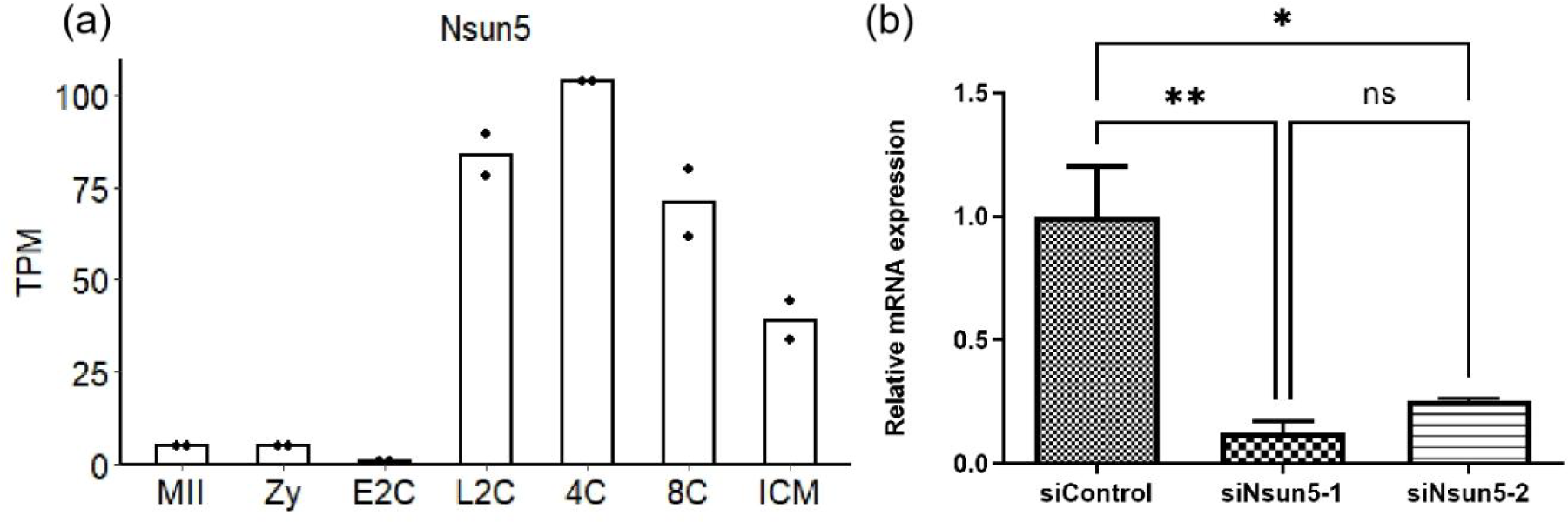
Expression profile of *Nsun5* mRNA in mouse preimplantation embryos and knockdown efficiency of siRNA treatment. **a** Expression profile of *Nsun5* mRNA during the mouse preimplantation development calculated from RNA-seq data (Zhang, et al. 2016). **b** Knockdown efficiency of siRNA treatment. All fertilized zygotes were microinjected with siControl or two different siRNAs targeting *Nsun5* within 3–6 hpi and subjected to RT-qPCR to access knockdown efficiency at 24 h after injection. Data are expressed as means ± standard error of the mean (S.E.M.) (three replicates). Ten to 15 embryos per replicate in each experimental group were analyzed. Statistical analysis was performed using one-way ANOVA followed by the Tukey-Kramer test. Not significant (ns) P > 0.1, *P < 0.05, **P < 0.01.

### *Nsun5* is essential for the development of mouse preimplantation embryos

We monitored embryo development from 24 to 120 hpi (Fig. 2). *Nsun5* knockdown significantly reduced the rate of mouse blastocyst formation (Fig. 2a and b). Remarkably, at 120 hpi, the blastocysts in the knockdown group exhibited a significant impairment in hatching from the zona pellucida when compared to the control group (Fig. 2a and c). Furthermore, at 96 hpi, the mean size of blastocysts was smaller in the knockdown group than in the control group (Fig. 2c and d). These results indicate the crucial role of *Nsun5* gene in regulating various aspects of early embryonic development, including blastocyst formation, hatching from the zona pellucida, and size development.

**Fig. 2.**
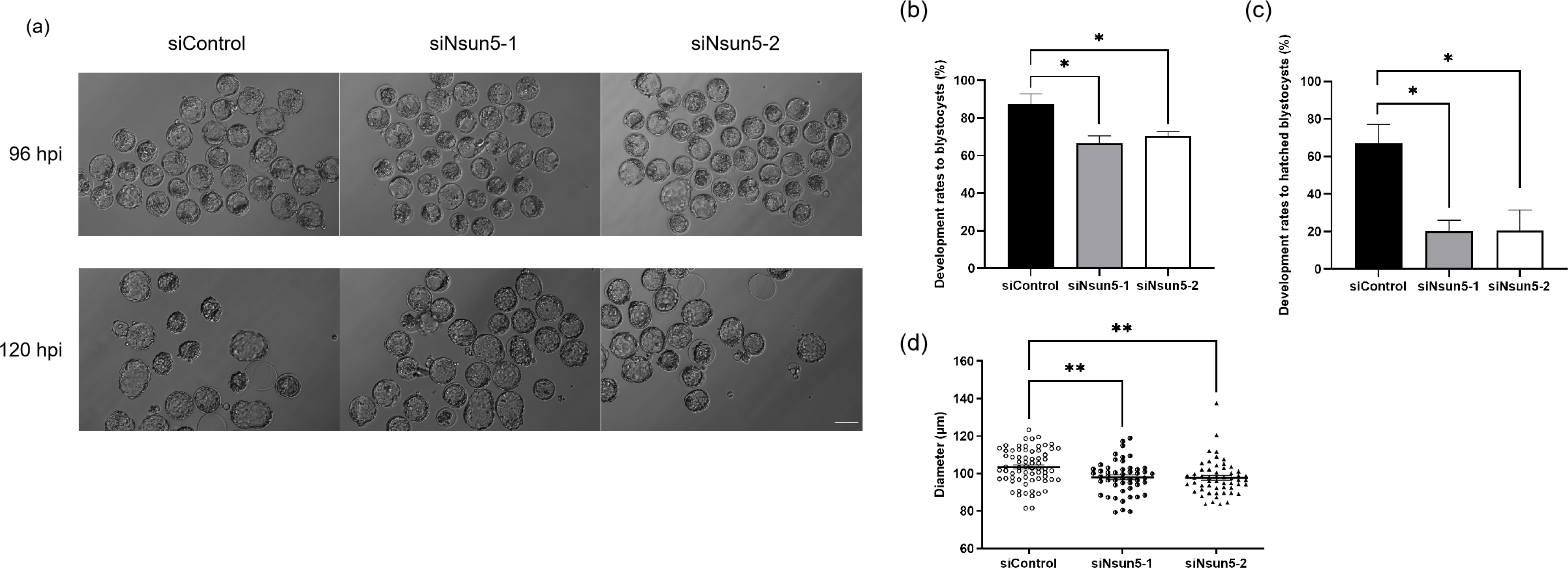
Effects of *Nsun5* knockdown on mouse preimplantation embryo development. **a** Morphology of mouse embryos after siControl or si*Nsun5* microinjection. Scale bar, 100 μm. Embryos were photographed at 96 and 120 hpi. **b** Blastocyst formation rate. **c** Hatched blastocyst rate. Zygotes were cultured *in vitro* until 120 hpi and the blastocyst rate and hatched blastocyst rate were quantified. Data are expressed as means ± S.E.M. (three replicates). siControl, *n* = 79; si*Nsun5*-1, *n* = 75; si*Nsun5*-2, *n* = 78. *P < 0.05, one-way ANOVA followed by the Tukey-Kramer test. **d** Diameter of blastocysts. siControl, *n* = 69; si*Nsun5*-1, *n* = 50; si*Nsun5*-2, *n* = 55. **P < 0.01, one-way ANOVA followed by the Tukey-Kramer test.

### *Nsun5* knockdown impairs cell proliferation in mouse preimplantation embryos

To investigate the underlying reasons for the impairment of embryonic development, we performed Hoechst staining of morula and blastocyst stage embryos to count cell numbers. At both stages, the *Nsun5* knockdown group exhibited a significant decrease in total cell number compared to the control group (Fig. 3a and b). These reductions suggest that *Nsun5* knockdown hampers cell proliferation during early embryonic development.

**Fig. 3.**
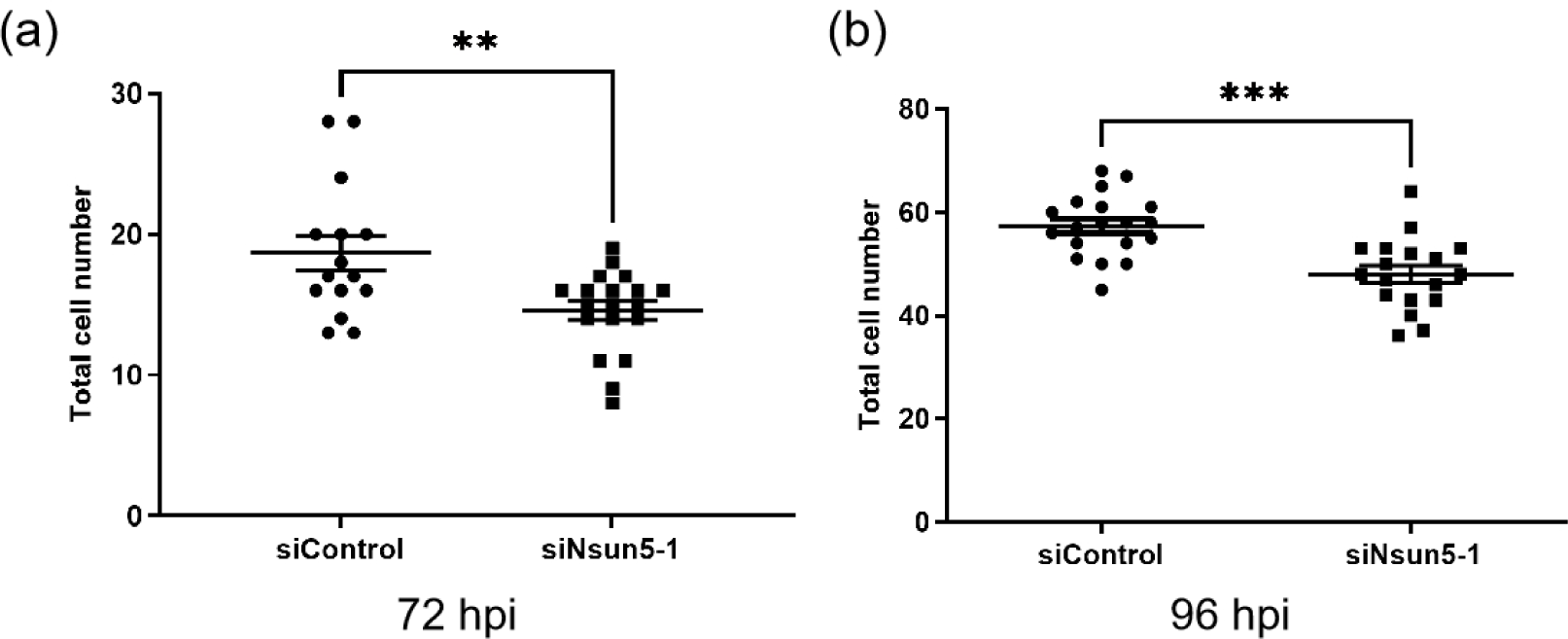
Effects of *Nsun5* knockdown on total cell number of embryos. **a** Total cell number of mouse morula stage embryos. siControl, *n* = 15; si*Nsun5*-1, *n* = 19. **P < 0.01, unpaired *t*-test. **b** Total cell number of mouse blastocyst stage embryos. siControl, *n* = 19; si*Nsun5*-1, *n* = 18. ***P < 0.001, unpaired *t*-test.

### *Nsun5* knockdown inhibits YAP1 nuclear translocation and impairs cell differentiation

As mentioned earlier, *Nsun5* depletion led to a severe consequence in which blastocysts were unable to hatch from the zona pellucida (Fig. 2b). This failure is possibly attributed to impaired cell differentiation and hindered development of trophectoderm cells (Wu, et al. 2010). The nuclear translocation of YAP1 during the morula stage is crucial for the formation of the trophectoderm (Rayon, et al. 2014). Therefore, to investigate the specific mechanisms underlying the inability of knockdown embryos to hatch from the zona pellucida, we performed YAP1 immunostaining on morula-stage embryos as well as CDX2 and OCT4 immunostaining on blastocyst-stage embryos.

We found that the nuclear translocation of YAP1 was inhibited at the morula stage in *Nsun5* knockdown embryos (Fig. 4a–c). Our data revealed a lower number of CDX2-positive cells in the blastocyst stage of the knockdown group (Fig. 4a, d). In addition, there was no difference in the number of OCT4-positive cells between the knockdown and control groups (Fig. 4a, f), while the knockdown group exhibited a significantly higher proportion of OCT4-positive cells in the total blastocyst cell population than the control group (Fig. 4a, g). Moreover, the ratio of OCT4-positive cells to CDX2-positive cells was also significantly higher in the knockdown group than in the control group (Fig. 4h).

**Fig. 4.**
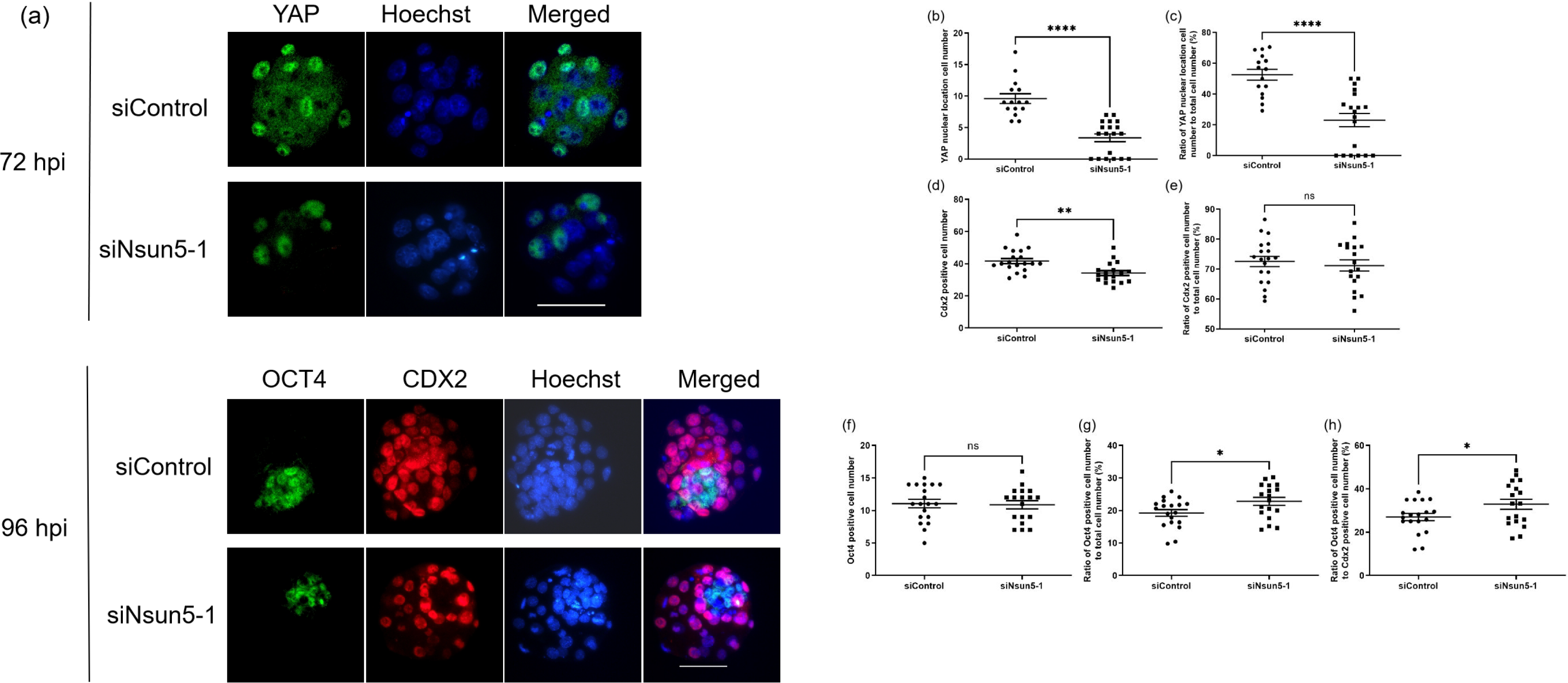
Effects of *Nsun5* knockdown on YAP1 nuclear translocation and cell differentiation of mouse preimplantation embryos. **a** Immunostaining images of mouse preimplantation embryos at the morula (72 hpi) and blastocyst (96 hpi) stages. Scale bar, 100 μm. **b** Number of cells that exhibited YAP1 nuclear localization at the morula stage. **c** Ratio of nuclear YAP1-positive cells to total cell number at the morula stage. siControl, *n* = 15; si*Nsun5*-1, *n* = 19. ****P < 0.0001, unpaired *t*-test. **d** CDX2-positive cells at the blastocyst stage. **e** Ratio of nuclear CDX2-positive cells to total cell number at the blastocyst stage. **f** OCT4-positive cells at the blastocyst stage. **g** Ratio of nuclear OCT4-positive cells to total cell number at the blastocyst stage. **h** Ratio of nuclear OCT4-positive cells to CDX2-positive cells at the blastocyst stage. siControl, *n* = 19; si*Nsun5*-1, *n* = 18. Ns P > 0.05, *P < 0.05, **P < 0.01, ****P < 0.0001, unpaired *t*-test.

### *Nsun5* knockdown alters the Hippo signaling pathway in mouse preimplantation embryos

To determine whether the impaired development observed in the *Nsun5* knockdown group was mediated through inhibition of the Hippo signaling pathway, we measured the relative mRNA levels of key genes involved in the Hippo signaling pathway at the morula stage, and assessed downstream genes controlled by the Hippo pathway that are associated with cell proliferation, cell differentiation, and hatching at the blastocyst stage. Our results demonstrated that *Nsun5* knockdown led to a significant increase in the relative mRNA levels of key genes in the Hippo signaling pathway, such as *Lats1* and *Lats2*, in the morula stage (Fig. 5a). Furthermore, the levels of *Mob1a* and *Mob1b*, which are also associated with the Hippo pathway, tended to increase in the *Nsun5* knockdown group (Fig. 5a). Additionally, at the blastocyst stage, *Nsun5* knockdown tended to increase the expression of *Notch1* (Fig. 5b), which is located downstream of the Hippo pathway and involved in hatching processes.

**Fig. 5.**
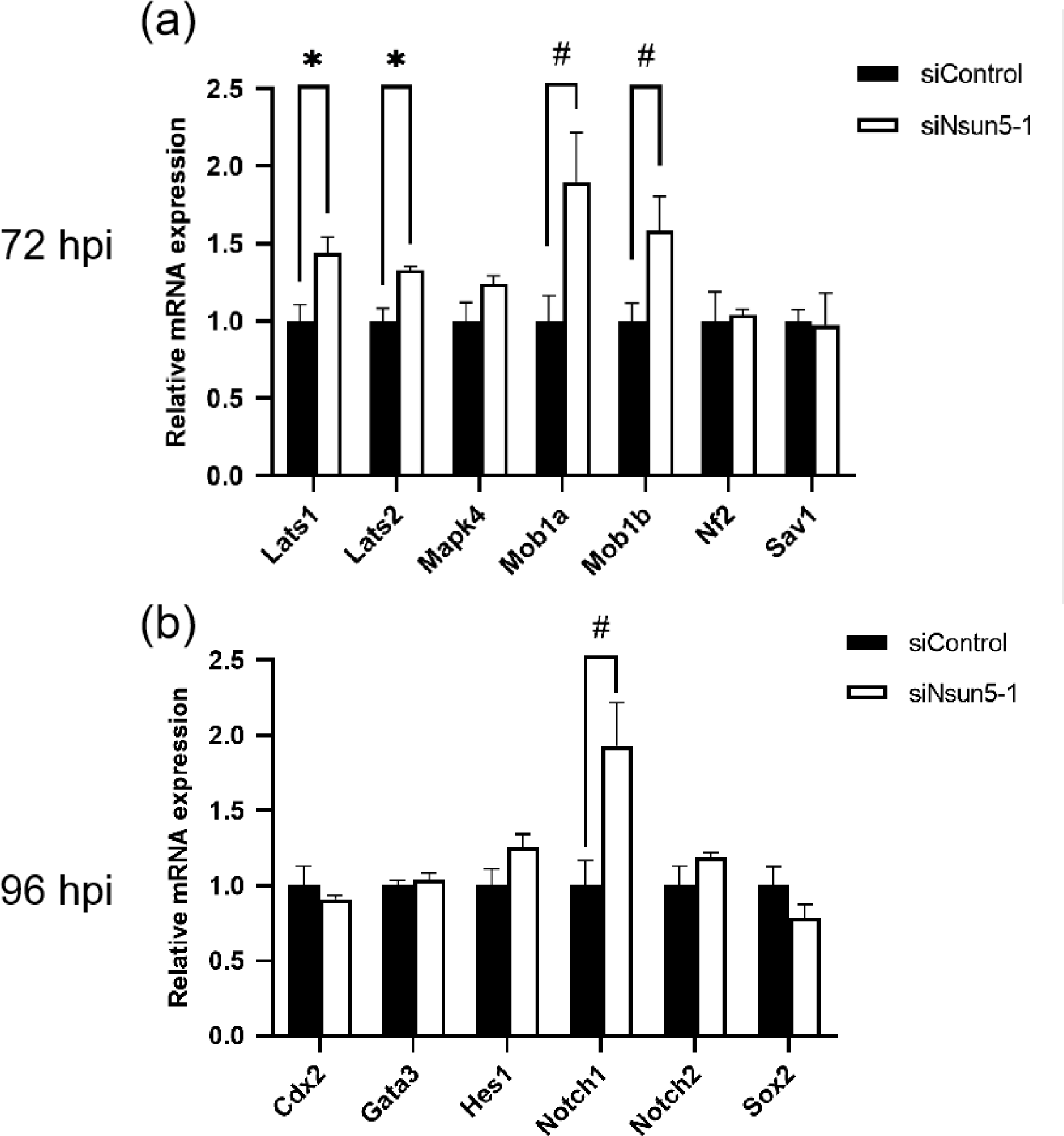
Expression levels of Hippo signaling-related genes at the morula and blastocyst stages. **A** Hippo signaling pathway genes were examined using morula stage embryos. **B** Hippo signaling downstream genes were examined using blastocyst stage embryos. Gene expression levels were normalized to *H2afz* as an internal control. Data are expressed as means ± S.E.M. (*n* = 3). Ten to 13 embryos per replicate in each experimental group were analyzed. #P < 0.1, *P < 0.05, unpaired *t*-test.

## Discussion

Previous studies suggest that RNA modifications play important roles in mouse preimplantation embryo development. The reversible N6-methyladenosine modification has vital functions in mammalian oocyte maturation and pre-implantation embryonic development processes (Sui, et al. 2020). N-acetyltransferase 10-mediated N4-acetylcytidine is a crucial regulator of oocyte maturation (Jiang, et al. 2023). m5C, one of the most prevalent modifications of RNA, is recruited by NSUN1–7 (Song, et al. 2022). Our study provides compelling evidence for the essential role of *Nsun5* in mouse preimplantation embryo development. By knocking down *Nsun5* mRNA levels, we observed impaired cell proliferation and differentiation and disrupted embryonic development, which aligns with findings from a previous study using *Nsun5* knockout mice (Ding, et al. 2022). The observed impairment in cell proliferation suggests that NSUN5 plays a crucial role in regulating cell cycle progression, consistent with a previous study demonstrating that decreased NSUN5 expression impairs cell cycle progression (Jiang, et al. 2020). Our findings shed light on a potential connection between *Nsun5* depletion, alterations in the Hippo signaling pathway, and the subsequent effects on YAP1 activity. *Nsun5* gene interference resulted in changes in the mRNA expression levels of key genes in the Hippo signaling pathway, such as *Lats1* and *Lats2*. LATS1 and LATS2 phosphorylate and inhibit nuclear translocation of the transcriptional co-activator YAP1 (Guo, et al. 2015, Zeng and Hong 2008). YAP1 plays a key role in cell fate determination (Yildirim, et al. 2021) and tissue development (Boopathy and Hong 2019), making it one of the most important downstream effectors of the Hippo signaling pathway (Genevet and Tapon 2011). The observed reduction in CDX2-positive cells during the blastocyst stage in the knockdown group suggests potential implications of NSUN5 for the inhibition of YAP1 nuclear translocation (Hirate, et al. 2015). This reduction may lead to the failure of knockdown embryos to form blastocysts (Fleming 1992) and results in limited expansion, which is essential for hatching and efficient interaction with the uterine endometrium during implantation in placental mammals (Marikawa, et al. 2012). This likely explains the challenges faced by knockdown embryos in hatching from the zona pellucida. Additionally, the altered proportions of OCT4-positive and CDX2-positive cells indicate a potential imbalance in the pluripotent and trophectoderm lineages in the knockdown embryos. These findings highlight the crucial role of NSUN5 in modulating YAP1 nuclear translocation, thereby influencing cell differentiation processes during preimplantation embryo development. Moreover, the observed developmental abnormalities in *Nsun5* knockdown embryos may be associated with dysregulation of the Hippo signaling pathway. The upregulation of key genes in the Hippo pathway and alterations in downstream gene expression provide insights into the molecular mechanisms underlying the developmental impairments observed in the knockdown embryos. However, more research is needed to reveal precisely how NSUN5 regulates these processes. Employing techniques such as RNA immunoprecipitation could uncover interacting partners and its role in Hippo pathway regulation.

In conclusion, our findings highlight the crucial role of *Nsun5* in mouse preimplantation embryo development. Knockdown of *Nsun5* mRNA led to impaired cell proliferation and differentiation, compromised YAP1 nuclear translocation, and alterations in the Hippo signaling pathway. The significance of *Nsun5*’s suggests potential therapeutic avenues for reproductive medicine and fertility.

## Author contribution statement

All authors conceived and designed the study. LD and SH performed experiments. All authors analyzed and interpreted the data. LD and SI drafted the manuscript. All authors discussed the results and reviewed, corrected, and approved the final manuscript.

